# Resolving human neuronal herpesvirus reactivation via petabase-scale association studies

**DOI:** 10.64898/2026.04.29.721769

**Authors:** Jacob C. Gutierrez, Yifan Chen, Artem Babaian, Ryan S. Dhindsa, Caleb A. Lareau

**Affiliations:** Computational and Systems Biology Program, Memorial Sloan Kettering Cancer Center, New York, NY, USA; Medical Scientist Training Program, Baylor College of Medicine, Houston, TX, USA; Jan and Dan Duncan Neurological Research Institute, Texas Children’s Hospital, Houston, TX, USA; The Donnelly Centre for Cellular & Biomolecular Research, University of Toronto, Toronto, Canada; Department of Molecular Genetics, University of Toronto, Toronto, Canada; Department of Pathology and Immunology, Baylor College of Medicine, Houston, TX, USA; Department of Molecular and Human Genetics, Baylor College of Medicine, Houston, TX, USA

## Abstract

Mounting evidence implicates herpesvirus reactivation in the etiology of Alzheimer’s disease, yet we lack a refined molecular characterization of pathogenesis in neurodegeneration. Here, we mine over 10 petabytes of human sequencing data for viral transcripts, identifying recurrent herpes simplex virus 1 (HSV-1) reactivation in healthy but not pathological post-mortem human brain tissue. Integrative single-nucleus analyses resolve direct evidence of HSV-1 expression in RORB+ glutamatergic neurons, implicating viral reactivation in a neuronal population progressively lost during dementia.

## MAIN

Although the etiology of Alzheimer’s Disease (AD) and related dementias involves a complex interplay of genetic predisposition^1^, metabolic dysfunction^2^, and vascular health^3^, a growing body of evidence implicates viral infection as another potential risk factor. Notably, a history of viral encephalitis has been linked to a substantially increased risk of AD,^4^ including from the neurotropic human herpes simplex virus 1 (HSV-1). Like all herpesviruses, HSV-1 can establish viral latency, wherein intact copies of the viral genome can persist for decades after primary infection^5^. HSV-1 can then reactivate from heterogeneous triggers, including infections by other viruses^6^ or following repeated mechanical injury akin to concussions^7^. Emerging evidence indicates that amyloid beta (Aβ) and phosphorylated tau (p-tau), hallmarks of AD pathology, neutralize HSV-1, suggesting that the pathology of AD may have evolved as an anti-viral innate immune response^8^. Despite decades of epidemiological and molecular evidence for a viral etiology of AD, direct transcriptomic evidence of HSV-1 reactivation in the human brain remains underdescribed, and no study has localized reactivation to specific cell populations.

Prior molecular studies on the role of HSV-1 in AD pathogenesis have relied largely on mouse models^9,10^ or *in vitro* systems. We therefore asked whether HSV-1 reactivation could be detected directly in existing human brain RNA-sequencing datasets. Because herpesvirus RNAs are polyadenylated during reactivation^11,12^, we reasoned that a systematic reprocessing of existing RNA sequencing data may resolve instances of HSV-1 reactivation in native human brain tissues, even when the virus was not originally noted. To characterize the landscape of HSV-1 reactivation in the context of AD as well as the general population, we reprocessed transcriptional profiling data from a combined 5,137 RNA-seq libraries curated through the Religious Orders Study and Memory and Aging Project (ROSMAP) and Genotype-Tissue Expression (GTEx) consortia (**Fig. 1a**). We developed a scalable pseudoalignment approach for HSV-1, similar to our previous efforts with EBV and HHV-6^11^, that showed limited false assignment to a viral reference (**Methods**). Application of this pipeline to these bulk RNA-seq samples resulted in 71,895,197 assigned HSV-1-derived transcripts from these two consortia, including four libraries from GTEx that exceeded 10 million viral reads (**Fig. 1b**).

**Figure 1.**
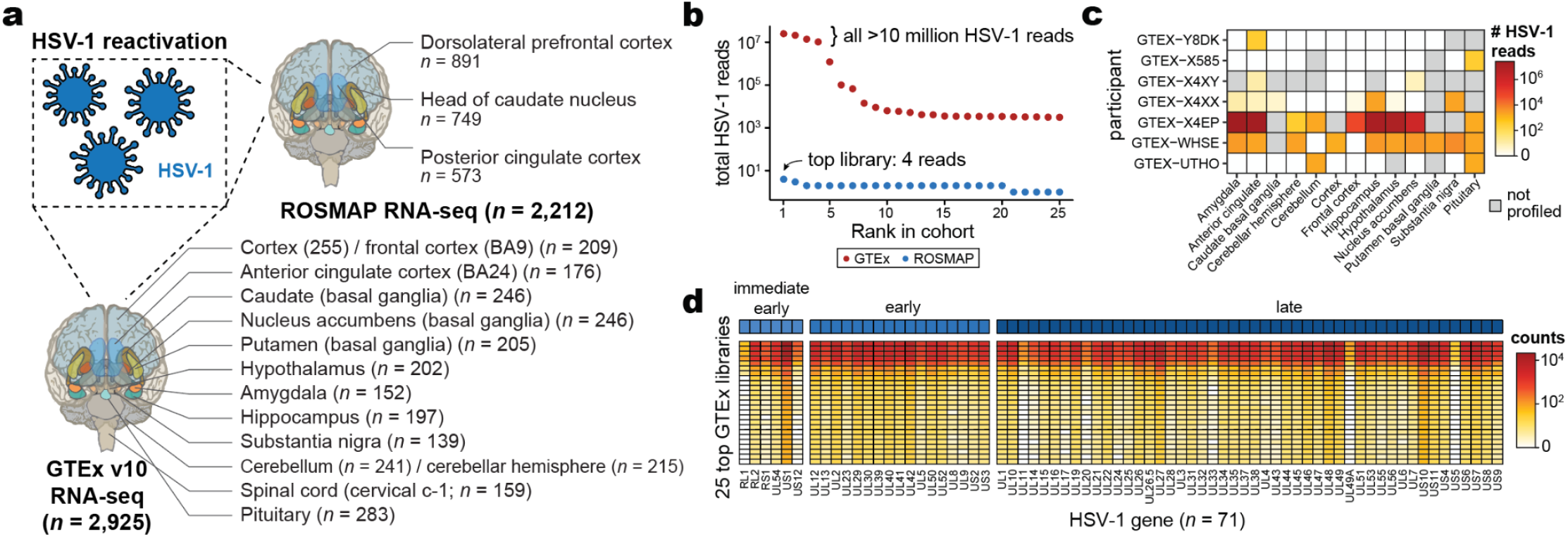
Assessment of HSV-1 latency and reactivation in native human tissues. **a**, Overview of data for inference of human herpes simplex virus 1 (HSV-1) across RNA-seq libraries from the Religious Orders Study and Memory and Aging Project (ROSMAP) and Genotype-Tissue Expression (GTEx) consortia. The *n* notes the number of libraries reprocessed. Schematic adapted from previous work^21^. **b**, Abundance of HSV-1 reads in the top 25 libraries per consortia. Annotations for the top 4 GTEx (all exceeding 10,000 HSV-1 reads) and top ROSMAP library (4 HSV-1 reads) are noted. **c**, Complete brain-associated data availability for 7 donors showing HSV-1 reactivation. **d**, Summary of HSV-1 expression at the gene level across 25 libraries with the highest HSV-1 expression.

Conversely, we observed minimal HSV-1 expression from ROSMAP samples, where the top library had only four assigned HSV-1 reads (**Fig. 1b**). Deeper analyses of the GTEx libraries showed seven distinct donors with HSV-1 reactivation, with the highest levels observed in one donor (GTEX-X4EP) in the amygdala, cingulate cortex, and hippocampus, confirming a prior report of reactivation in this donor from a distinct bioinformatic pipeline^13^ (**Fig. 1c**). Further, we noted that the HSV-1-positive libraries were processed in eight independent batches, strongly suggesting that these reads did not result from contamination across library construction or sequencing (**Extended Data Fig. 1a**). Across all libraries, we observed 71 total HSV-1 genes expressed that spanned the entire viral life cycle, verifying true viral reactivation rather than latency (**Fig. 1d**). HSV-1 reactivation is therefore detectable across multiple brain regions in the general population but is essentially absent from ROSMAP, a post-mortem cohort enriched for older individuals for AD neuropathology.

Encouraged by our ability to detect HSV-1 reactivation in GTEx and ROSMAP, we next scaled our analysis to the Sequence Read Archive (SRA) using Logan, querying 11.9 petabases of human sequencing data, irrespective of library design (**Fig. 2a; Methods**). In brief, we used the LoganSearch interface, which contains a 31-mer index of all unitigs and contigs assembled from the entirety of the SRA (**Fig. 2b**). As queries, we selected the three shortest HSV-1 protein-coding genes in the reference genome, noting all had direct homologues with other species from the *Herpesviridae* family, including Varicella Zoster Virus (VZV/HHV-3; **Extended Data Fig. 2a; Methods**). The result of these queries were libraries with human cells and nuclei as input to library preparation and when one or more high-confidence HSV-1-derived contigs were detected, reflecting viral infection, latency, and/or reactivation (**Fig. 2c**). Over the >1,000 libraries, most results were instances where HSV-1 was deliberately introduced into human cells to study host responses, including infection of primary skin fibroblasts^14^ and cerebral organoids^15^. These studies served as positive controls for the workflow. However, to identify instances of true reactivation rather than experimental infection, we focused on libraries in which HSV-1 expression was not annotated in the original study, building on a prior survey of HHV-6 reactivation in therapeutic T cell cultures^11^.

**Figure 2.**
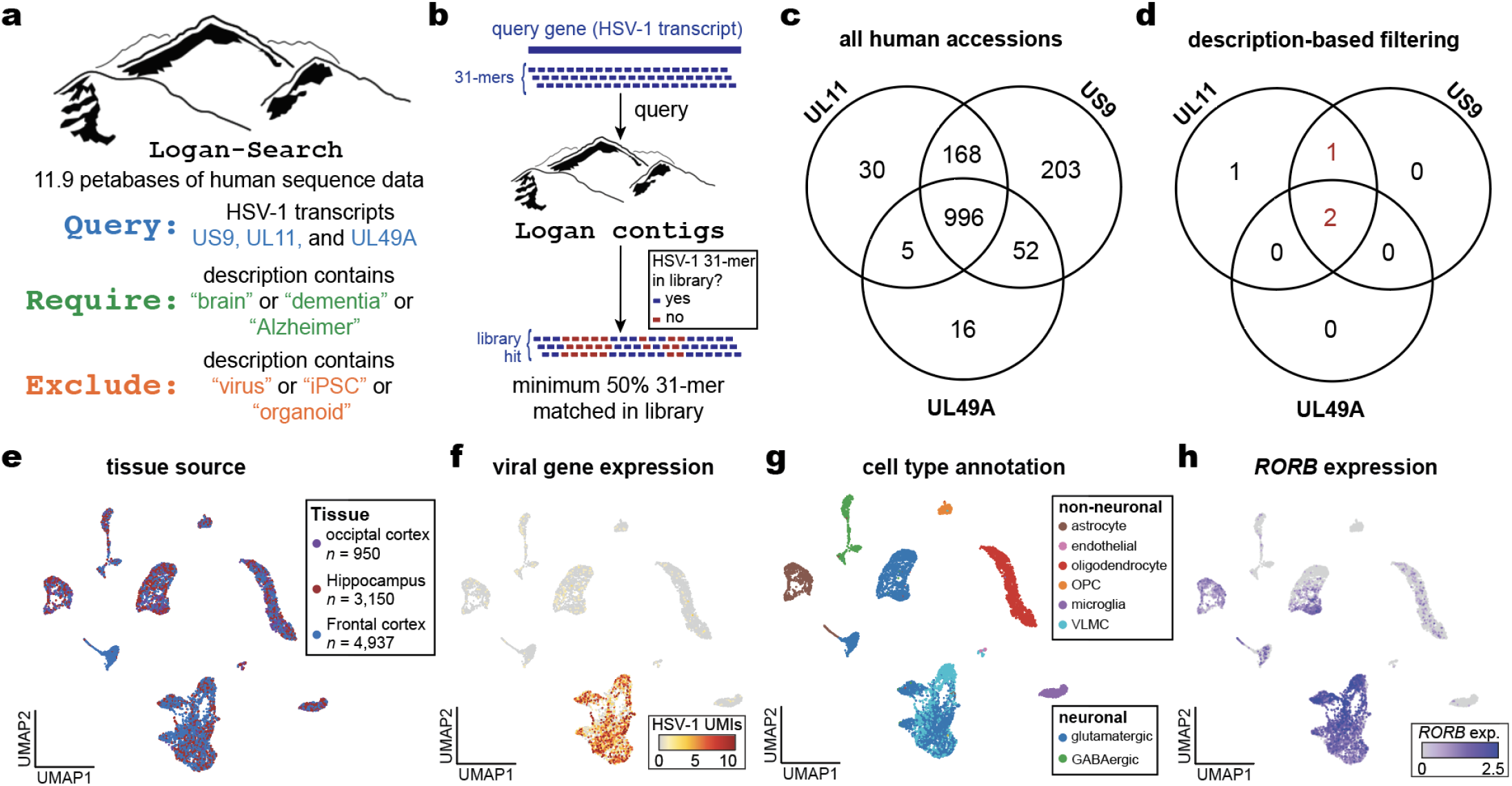
Resolving HSV-1 reactivation in RORB+ glutamatergic neurons. **a**, Schematic of Logan-Search input queries and downstream filtering criteria. **b**, Schematic of LoganSearch matching of HSV-1-associated 31-mers. **c**, Summary of human input libraries with detected HSV-1 expression from one or more input genes. The Venn Diagram enumerates libraries with 50% of 31-mers in the query gene detected in the Logan compression of the accession. **d**, Count of libraries following text description filtering (“Require” and “Exclude” criteria in **a**). **e**, Uniform Manifold Approximation and Projection (UMAP) embedding of scRNA-seq libraries with HSV-1 detected from **d. f**, Per-cell quantification of total HSV-1 expression. **g**, Cell type annotation. **h.** *RORB* expression in the same cells.

To isolate potential brain-associated viral reactivation, we applied text-based filtering to the metadata descriptions of the accession. Our heuristics i) excluded libraries where the description corresponded to viral, iPSC, or organoid studies, and ii) retained sequencing datasets that mentioned “brain”, “dementia”, or “Alzheimers” (**Fig. 2a**; **Methods**). Three sequencing datasets detected by two or more queries were all from the same individual study (**Fig. 2d**)^30^. These three accessions mapped to a single study of single-nucleus RNA-sequencing libraries collected from three brain regions of a single individual. The profiled individual was a carrier of the *PSEN1* E280A mutation, the most common cause of familial early-onset Alzheimer’s disease.

Notably, she was protected against Alzheimer’s symptoms for almost three decades beyond the expected age of onset, and her resilience was attributed to homozygosity for the *APOE3* Christchurch (R136S) variant^30^.

Despite a very high amyloid burden, tau pathology and neurodegeneration in her brain were largely confined to the occipital cortex, with the frontal cortex and hippocampus relatively spared. These regions retained remarkable populations of RORB+ excitatory neurons, a selectively vulnerable population typically depleted in AD^16–18^.

To assess the tropism of the putative reactivated HSV-1, we reprocessed the raw single-nuclei data derived from the occipital cortex, hippocampus, and frontal cortex. Notably, as transcripts from the *Herpesviridae* family are poly-adenylated^11^, mapping HSV-1 expression could refine the specific cell populations harboring reactivated HSV-1. Reprocessing of reads to a joint human-HSV-1 reference revealed that HSV-1 expression was predominantly concentrated in the hippocampus (**Table 1a; Fig. 2e,f**).

**Table 1.** Summary of herpesvirus expression in single-cell libraries. **a**, Quantification of herpesvirus transcripts from the HSV-1 reactivation libraries nominated via Logan^25^ and **b**, Summary of additional scRNA-seq library quantifications from atlases of cells and tissues profiled from AD donors.

| **a. Description** | **Accession** | **# cells / nuclei** | **# donors** | **# raw reads** | **#HSV-1** |
| --- | --- | --- | --- | --- | --- |
| scRNA-seq library of occipital cortex | SRR19792154 | 1,423 | 1 | 46,789,272 | 835 |
| scRNA-seq library of hippocampus | SRR19792155 | 5,751 | 1 | 159,833,749 | 1,500,373 |
| scRNA-seq library of frontal cortex | SRR19792156 | 7,712 | 1 | 165,740,616 | 95,078 |
| **b. Description** | **Accession** | **# cells / nuclei** | **# donors** | **# raw reads** | **#HSV-1** |
| Prefrontal cortex atlas of AD patients<sup>22</sup> | syn52293417 | 2,359,994 | 427 | 179,638,739,333 | 0 |
| Frontal cortex atlas of AD patients<sup>23</sup> | syn16780177 | 172,659 | 24 | 2,834,676,662 | 0 |
| Prefrontal cortex atlas of AD patients<sup>24</sup> | syn18485175 | 80,660 | 48 | 5,375,838,165 | 0 |

Dimensionality reduction, clustering, and cell type reference projection of the nuclei profiles localized the signal to a specific population of glutamatergic neurons, including 105 HSV-1 super-expressor cells^11^ (**Fig. 2g; Extended Data Fig. 2b,c**). Notably, this population is marked by *RORB* expression, a transcription factor that underlies specific neuronal functional phenotypes^18^ (**Fig. 2h**). We evaluated whether the HSV-1 reactivation in this donor could be explained by an attenuated form of the virus, however we detected no deleterious variants among high-confidence mutations in the viral genome (**Extended Data Fig.2d)**. Together, our reanalysis of these libraries suggests that this RORB+ glutamatergic neuron population progressively lost in AD^18–20^ may be susceptible due to viral reactivation and degeneration as an early event in initially healthy individuals that ultimately develop dementia.

To further assess HSV-1 reactivation in neuronal populations, we downloaded and reanalyzed transcriptional profiles from >2.6 million nuclei derived from 499 donors with AD (**Table 1b**). Despite this collection of samples spanning far more donors than the Christchurch case identified by Logan, we did not detect a single high-confidence HSV-1-derived UMI from >185 billion reads. This null result extends our observation of the lack of HSV-1 expression from our bulk profiles in GTEx and ROSMAP, in which donors with AD neuropathology had no detectable viral transcripts. The one brain in which we detected HSV-1 reactivation is also the one brain in which the vulnerable RORB+ neurons had been preserved. We interpret this pattern as evidence of survivor bias: the neurons that reactivate HSV-1 in AD are cleared before autopsy, rendering viral transcripts undetectable in standard post-mortem cohorts and detectable only in tissue protected from the neuronal loss that defines the disease.

Taken together, our data support a model in which RORB+ neurons in the hippocampus and frontal cortex serve as reservoirs of latent, ultimately reactivated HSV-1. Because these neurons are selectively vulnerable in AD^18–20^, they are likely lost following reactivation, accounting for their depletion and for the absence of detectable HSV-1 transcripts in post-mortem AD brain samples. While further work is needed to identify suitable interventions for this phenomenon, our study reinforces the immense value of comprehensive annotation of unmapped sequencing reads in resolving novel host-viral interactions that may underlie complex human disease.

## EXTENDED DATA FIGURES

**Extended Data Fig. 1.**
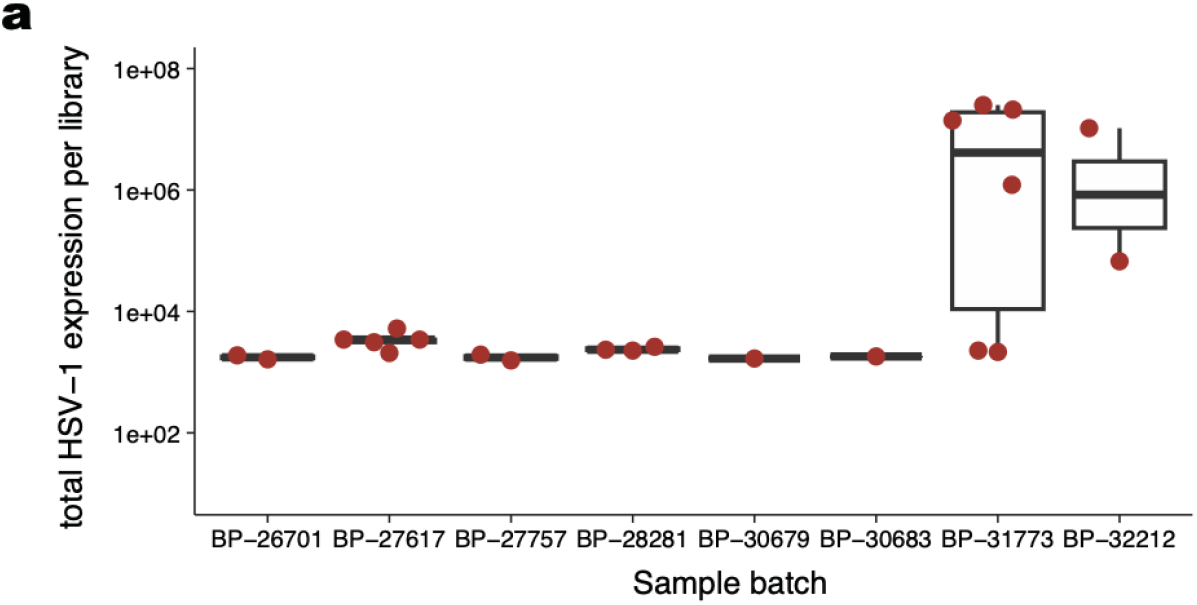
Supporting information for GTEx reanalyses. **a**, Summary of HSV-1 detection across eight processing batches.

**Extended Data Fig. 2.**
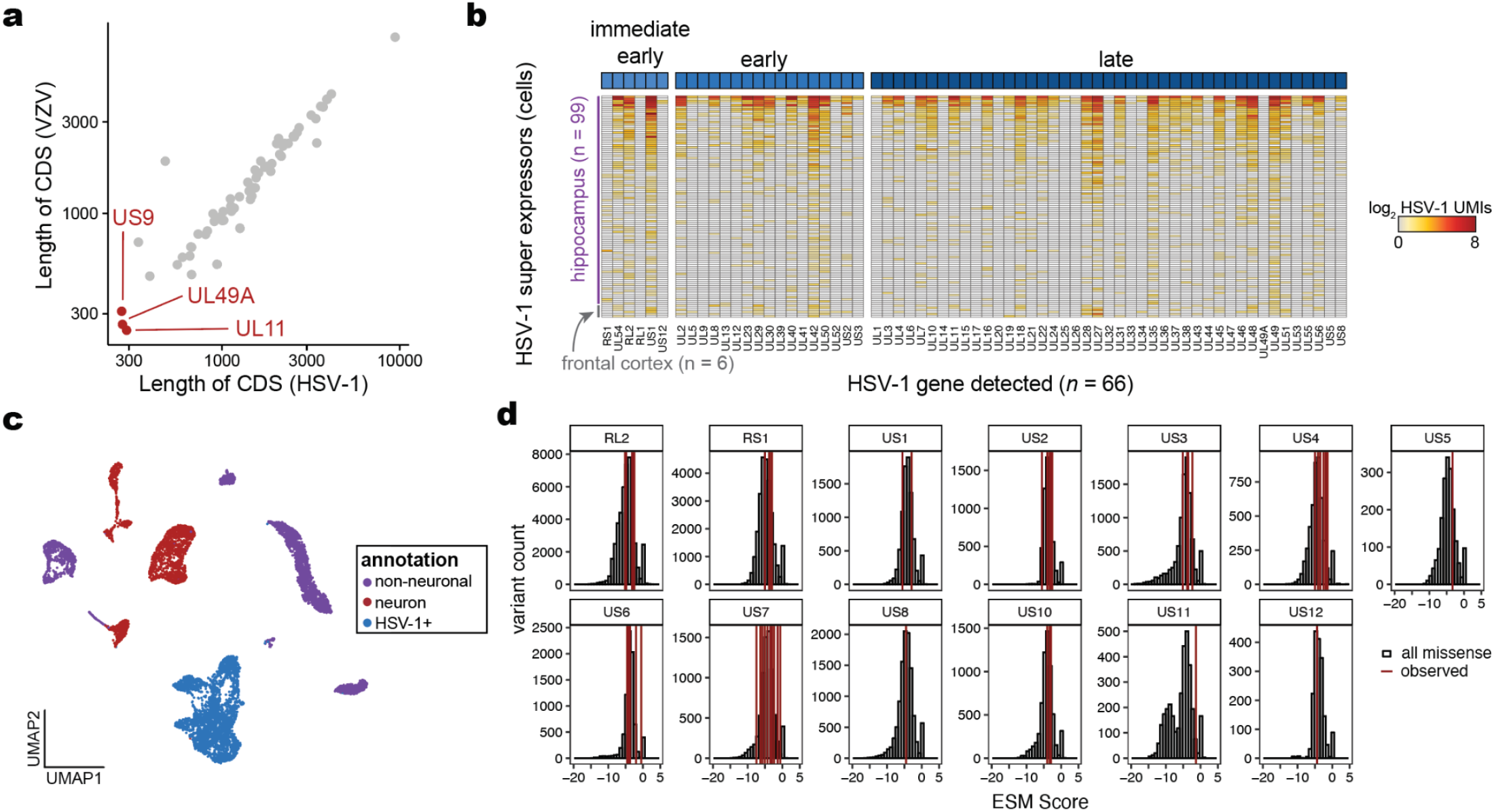
Supporting information for Logan and scRNA-seq analyses. **a**, Rationale of selected herpesvirus coding DNA sequences (CDS) in HSV-1 (*UL11, UL49A*, and *US9* genes) based on contig length and conservation across other *Herpesviridae* species. **b**, Characterization of transcripts expressed from 105 HSV-1 super-expressors (minimum 10 viral-derived UMIs) from scRNA-seq. **c**, Annotation of populations from single-cell profiles. **d**, Characterization of HSV-1 strain mutations identified in donor via ESM^26^.

## METHODS

### HSV-1 reference genome

The primary reference genome for this study was NC_001806.2 (HSV-1), which was curated through GenBank. As species from the *Herpesviridae* family mutate relatively slowly compared to other viral families^27^, our analyses could proceed using only a singular reference genome, both for seeding queries in Logan and reanalyzing existing RNA-seq datasets. Applications to other viral families characterized by more rapid evolution would require the use of a more extensive viral pangenome.

### GTEx and ROSMAP reanalyses

Raw .fastq reads were steamed from the ROSMAP synapse instance using the synapseclient PyPi package. For GTEx, unmapped RNA-seq reads were accessed on Terra.bio and streamed via samtools view^28^. We noted that the GTEx RNAseq libraries were prepared with poly-A tail enrichment, while the ROSMAP cohort generated libraries by rRNA depletion. We noted this as rDNA depletion should have higher sensitivity to detect viral reads, reinforcing the interpretation of **Fig. 1b** that the variable presence of the virus was driven more by disease stage than a specific technical variable. For both cohorts, reads were quantified using a kallisto index of the NC_001806.2 coding sequence (CDS) contig indexed with 31-mers and subsequently analyzed using the abundance.tsv file coming from the ‘quant’ mode.

For the GTEx cohort, donors who were positive for HSV-1 reactivation were not significantly different among any obvious covariates, including donor sex (*p* = 0.44; two-sided Fisher’s Exact Test) or age (*p* = 0.57; two-sided Student’s *t*-test). Five unique causes of death were attributed to the donors with reactivated brain HSV-1, including heart disease, blunt force trauma from a motor vehicle accident, cerebrovascular accident, and cardiac arrest. Additional metadata, including sequencing batch, was analyzed for any interpretable confounder that would explain the specificity of HSV-1 expression, but no obvious expression signature could be identified (**Extended Data Fig. 1a**).

### LoganSearch Queries

LoganSearch is a free-to-use, web interface that allows for queries of up to 1 kilobase and returns accessions with a minimum of 25% of the 31-mers in the query sequence found in the assembled unitigs and contigs^29^. We queried LoganSearch on January 5, 2026 (v0.8.0) using the web interface and supplied each of the HSV-1 genes noted in **Extended Data Fig. 2a** in three separate queries. The output of the queries was downloaded locally in .csv format and processed using a custom R script, where the associated ‘description’ metadata was filtered using case-insensitive ‘grep’ searches.

### Single-cell RNA-seq reanalyses

After identifying the accessions from the LoganSearch query, we downloaded raw .fastq files from the SRR accessions and requantified using CellRanger v8.0.1. We aligned these reads using the GRCh38 2024-A GenCode.v44 GTF, which was modified to include the HSV-1 sequence (NC_001806.2) using ‘cellranger mkref’ command with default parameters. We note that this additional step of re-mapping associations to a reference genome is required to infer specific viral genes that are expressed in these samples as Logan does not label viral genes. In our experience reanalyzing existing scRNA-seq data for herpesvirus expression^11^, true-positive identification of viral reactivation constitutes multiple genes being expressed, whereas only a single gene is more prone to erroneous read mapping artifacts.

To mitigate the impact of ambient RNA, the filtered cellcount .h5 files were used as input to SoupX v1.6.2^30^, which resolves high-confidence HSV expression associated with cell droplets. These corrected matrices were then used for doublet detection using scDblFinder v1.18.0 ^31^ followed by downstream analysis in Seurat v5.4.0 ^32^. We split these expression matrices by human and HSV-1 genes into separate assays to ensure HSV expression did not contribute to cell state analysis. Similar to the source publication ^25^, cells were filtered with the following subset command: ‘nFeature_RNA > 200 & percent.mt < 5 & percent.ribo < 10’.

SCTransform followed by CCA integration with default parameters was applied to create a joint representation of all samples. UMAP embeddings and clustering were created using 20 PCs from the integrated PCA for all downstream analysis. High-confidence HSV+ cells were annotated by the detection of at least 3 HSV UMIs due to the prevalence of cells with 1 or 2 HSV reads. We defined 105 HSV super-expressor cells using our same definition of a minimum of 10 ambient-correct viral UMIs per high-quality cell, akin to our definition of HHV-6 super-expressor cells^11^. Though other clusters had non-zero HSV-1 read abundance, we caution against interpreting tropism or reactivation from these data due to ambient viral RNA that can cross-contaminate during 10x Genomics library preparation^10^. Metadata for single cells was downloaded from the source GEO link and joined to the object to understand how previous work interpreted these cell states. To label transfer brain cell types onto our dataset, we used Azimuth v0.5.0 with the motor cortex reference.

We reasoned that one possibility for viral expression in this individual’s brain could be an attenuated form of the HSV-1 virus that resulted in a less pathogenic expression. To assess this, we conducted analyses of viral transcripts for potential mutations by extracting only the HSV-1 aligned reads via samtools view. These HSV-1-specific .bam files were used as input to freebayes v1.3.6 ^33^ to identify the viral variants present in the combined donor viral reads. The resulting .vcf files were then annotated with VEP v113.0 using the HSV-1 .gff file downloaded from GenBank. SNVs of interest were kept if they were concordant in both of the SRR19792155 and SRR19792156 samples. This resulted in a total of 68 high-confidence HSV-1 derived variants with predicted protein-coding changes from VEP. To interpret these variants, we used ESM predictions^26^ from all possible mutations in the full HSV-1 protein .fasta file to assess all possible amino acid alterations and their ESM score of predicted consequence. This analysis revealed no outliers in loss-of-function alleles in any variants observed in the HSV-1 genome, which we interpret as no clear evidence that the viral strain was attenuated (**Extended Data Fig. 2d**).

Finally, for additional scRNA-seq analyses (**Table 1b**), selected AD studies were identified based on data availability in Synapse and streamed locally for downstream analyses. Presence of HSV-1 reads was assessed using kallisto with the same viral genome file as the index. Due to the lack of viral reads being detected, no downstream single-cell analyses were performed. We note that each AD study did not include a typical case/control design; instead, donors were scored for AD pathology and cognitive impairment along a spectrum capturing a full breadth of brain states.

## Acknowledgements

We thank members of the Babaian, Dhindsa, and Lareau labs for helpful discussion. We acknowledge R. Chikhi for support with Logan analyses. This work was directly supported by an American Brain Foundation Inflammation award (A.B., R.S.D., C.A.L.), NIH grants P30CA008748 (C.A.L.), U01AT012984 (C.A.L.), a Michelson Prize Next-Generation Grant (C.A.L.), and a National Academy of Medicine Catalyst Prize (C.A.L.).

## Code availability

Custom code for analyses is available online at https://github.com/clareaulab/ad-hsv-mapping.

## Data availability

No new sequencing data were generated as part of this work. Raw GTEx reads are accessible via dbGaP and were accessed via an MSK application (#37709). LoganSearch is freely available at https://logan-search.org/. Single-cell profiles of Alzheimer’s Disease nuclei are available on synapse.org and accessed under an MSK application (#9603055).

